# GPU-Accelerated Optimization Investigates Synaptic Reorganization Underlying Pathological Beta Oscillations in a Basal Ganglia Network Model

**DOI:** 10.64898/2026.04.16.718939

**Authors:** Kavin Ranak Nakkeeran, William S. Anderson

## Abstract

**Objective:** Pathological beta-band oscillations (13 to 30 Hz) in the subthalamic nucleus (STN) are a hallmark of Parkinson’s disease and a primary target for deep brain stimulation therapy, yet the specific pattern of synaptic reorganization that drives their emergence remains incompletely understood. We developed a GPU-accelerated computational framework to systematically investigate combinations of synaptic changes across basal ganglia pathways that produce Parkinsonian beta oscillations while satisfying literature-based electrophysiology constraints.

**Approach:** We implemented a biophysically detailed spiking network model of the STN, external globus pallidus (GPe), and internal globus pallidus (GPi) in JAX (a high-performance numerical computing Python library), achieving a 490-fold speedup over conventional CPU-based simulation. Using the Covariance Matrix Adaptation Evolution Strategy (CMA-ES) we optimized 10 network parameters across two stages: first establishing a healthy baseline matching primate electrophysiology data, then searching within biologically motivated bounds for synaptic modifications that reproduce Parkinsonian firing rates and beta power. Fixed in-degree connectivity ensured optimized parameters produced scale-invariant dynamics from 450 to 45000 neurons. All simulations ran on a single cloud GPU instance at 84 cents per hour.

**Main Results:** The optimizer converged on a coordinated pattern of synaptic reorganization dominated by asymmetric changes within the STN-GPe reciprocal loop: STN to GPe excitation increased 2.21-fold while GPe to STN inhibition collapsed to 0.11-fold of its healthy value. STN to GPi and GPe to GPi pathways changed minimally (1.06-fold and 1.45-fold respectively). This configuration transformed asynchronous firing (beta: 0.4 percent of spectral power) into synchronized bursting with prominent beta oscillations (49.4 percent), with firing rate changes matching experimental observations. Network dynamics were invariant across a 100-fold range of network sizes (firing rate deviation less than 2.4 Hz; all metrics p less than 0.001 across 10 random seeds at 45000 neurons). We implemented a simplified deep brain stimulation model for validation purposes, which achieved complete beta suppression (49.4 percent to 0.0 percent) and restored GPi output to healthy levels.

**Significance:** These results suggest that pathological beta oscillations emerge from a specific pattern of synaptic reorganization, namely the reduction of GPe inhibitory feedback to STN. The GPU-accelerated optimization framework, running on commodity cloud infrastructure, demonstrates an accessible platform for parameter exploration in neural circuit models and a foundation for generating synthetic training data for adaptive deep brain stimulation algorithms.

## 1. Introduction

Parkinson’s disease (PD), the second most prevalent neurodegenerative disorder, affects over 10 million individuals worldwide, and is characterized clinically by tremor, rigidity, bradykinesia, and postural instability [1]. The physiological basis of these motor symptoms has been traced to disruptions in basal ganglia circuit dynamics following the loss of dopaminergic neurons in the substantia nigra pars compacta. A critical advance in understanding these disruptions came from the discovery that Parkinsonian motor symptoms are accompanied by abnormally sustained in the beta band oscillatory activity (13–30 Hz), recorded from the subthalamic nucleus (STN), globus pallidus, and cortex of both PD patients and animal models [2–4]. Beta power correlates with motor symptom severity [5], is reduced by dopaminergic medication [6], and is suppressed by clinically effective deep brain stimulation (DBS) of the STN [7, 8]. The causal relationship between beta oscillations and motor impairment has motivated their use as a control signal for adaptive, closed-loop DBS systems [9, 10].

Despite the clinical significance of beta oscillations, the exact synaptic mechanisms through which they emerge remain incompletely characterized. Experimental studies using electrophysiological recordings in 1-methyl-4-phenyl-1,2,3,6-tetrahydropyridine (MPTP)-treated primates have documented elevated and bursty STN firing, reduced and irregular globus pallidus externa (GPe) activity, and altered pallidal output to thalamus [11–13]. At the circuit level, the reciprocal excitatory-inhibitory loop between STN and GPe has long been identified as a candidate pacemaker for beta oscillations [14, 15]. Computational models have demonstrated that this loop can sustain oscillatory activity when the balance between excitation and inhibition is disrupted [16, 17]. Here, we investigate which specific pattern of synaptic reorganization, among the many changes occurring simultaneously under dopamine depletion, is responsible for destabilizing this circuit and driving the transition from healthy to pathological dynamics.

Addressing this question computationally has been limited by a practical bottleneck. Biophysically detailed models of the STN-GPe-GPi network require numerical integration of thousands of Hodgkin-Huxley neurons with voltage-dependent synaptic conductances at sub-millisecond timesteps, making individual simulations computationally expensive. Systematically searching a high-dimensional parameter space across synaptic conductances, intrinsic currents, and noise levels requires thousands of such simulations. This is restrictive on conventional CPU hardware as a single 500 ms simulation of even a modest 450-neuron network requires approximately 14 minutes using standard NumPy implementations. Prior modeling studies have relied on either simplified neuron models (leaky integrate-and-fire, firing rate models) that sacrifice biophysical realism [14, 16], manual parameter tuning [17, 18], or small networks that may not reproduce emergent population dynamics [19]. These constraints have meant that the synaptic basis of beta oscillations has been explored through perturbation of individual parameters rather than data-driven discovery.

Here, we overcome this computational bottleneck by implementing the STN-GPe-GPi network model in JAX [20], a high-performance numerical computing framework that enables just-in-time (JIT) compilation of Python code to optimized GPU kernels via the XLA compiler. Deployed on cloud computing infrastructure (Google Cloud Platform), this implementation achieves a 490-fold speedup over conventional CPU-based simulation, reducing the wall-clock time for a 450-neuron, 600 ms simulation from 837.75 seconds to 1.71 seconds on a single NVIDIA L4 GPU. The entire pipeline requires no local GPU hardware, running on commodity cloud instances accessible to any laboratory with an internet connection.

This platform enabled a complete 1,000-trial optimization of 10 network parameters, six intrinsic current and noise parameters plus four synaptic conductance multipliers, in under one hour. We employed Covariance Matrix Adaptation Evolution Strategy (CMA-ES) [21] within the Optuna optimization framework [22] to search for synaptic configurations that reproduce healthy and Parkinsonian firing statistics derived from the primate electrophysiology literature [11–13]. The optimizer independently discovered a coordinated pattern of synaptic reorganization between the healthy and the Parkinsonian states. STN→GPe excitation increased 2.21x, GPe→STN inhibition collapsed to 0.11× its healthy value, STN→GPi excitation remained near baseline (1.06×), and GPe→GPi inhibition increased 1.45x. This mechanism, where excitatory drive escalates while the inhibitory brake on STN fails, emerged from the optimization within biologically motivated search bounds that reflected known directions of dopamine-dependent plasticity (see Methods Section 2.7 for an account of the search range constraints). The resulting network exhibits robust beta-band activity (49.4% of STN spectral power in 13–30 Hz, compared to 0.4% in the healthy state), dynamics that are invariant across a 100-fold range of network sizes (450 to 45,000 neurons), and complete beta suppression under simulated high-frequency DBS.

These results provide three contributions. First, they suggest a specific pattern of synaptic reorganization as a candidate mechanistic basis for pathological beta oscillations, consistent with known patterns of dopamine-dependent plasticity. Second, they demonstrate that GPU-accelerated optimization can discover biologically plausible circuit mechanisms without manual parameter tuning, enabling a data-driven approach to computational neuroscience. Third, they establish an accessible, cloud-based computational platform that any laboratory can deploy to systematically explore neural circuit dynamics at scale.

## 2. Methods

### 2.1 Network Architecture

The model consists of three interconnected populations representing the subthalamic nucleus (STN), external segment of the globus pallidus (GPe), and internal segment of the globus pallidus (GPi), maintained in a 2:4:3 ratio. The optimization procedure was conducted at a network size of 450 neurons (100 STN, 200 GPe, 150 GPi), and validation simulations scale to 45,000 neurons (10,000 STN, 20,000 GPe, 15,000 GPi).

Connectivity follows a fixed-indegree scheme in which each postsynaptic neuron receives a constant number of presynaptic inputs *K*, sampled uniformly without replacement from the presynaptic population. This approach, recommended by Gerstner et al. [23] for network models intended to generalize across sizes, preserves both the mean and variance of total synaptic input to each neuron regardless of population size. In contrast, the more common fixed connection probability approach preserves only the mean, while its variance scales as 1/√*N*, progressively suppressing fluctuation-driven dynamics in larger networks. The indegree values were derived from connection probabilities established at the 450-neuron optimization size: STN→GPe *K* = 15, GPe→STN *K* = 14, STN→GPi *K* = 30, GPe→GPi *K* = 10 (Table 1). This architecture enables efficient parameter discovery at small, computationally tractable network sizes, with optimized parameters transferring directly to larger networks without retuning. Structural verification confirmed that every postsynaptic neuron receives exactly the prescribed number of inputs at all network sizes tested (450–45,000 neurons).

Four synaptic pathways connect the populations. STN→GPe and STN→GPi are excitatory (glutamatergic, AMPA-mediated), while GPe→STN and GPe→GPi are inhibitory (GABAergic, GABA_A-mediated). The architecture does not explicitly model cortical input, striatal projections, or thalamic feedback, rather, these afferent influences are approximated by the tonic drive currents and stochastic background input to each population (see Sections 2.2 and 2.4). The implications of this simplification are addressed in the Discussion.

### 2.2 Neuron Models

#### 2.2.1 STN Neurons

STN neurons are modeled as single-compartment Hodgkin-Huxley neurons based on the characterization by Gillies and Willshaw [17]. The membrane potential evolves according to C_m dV/dt = −I_Na − I_K − I_L − I_T − I_CaH − I_AHP − I_H − I_syn + I_drive + I_ext, where C_m = 1.0 µF/cm². The model includes seven ionic currents whose parameters are listed in Table 1a. The T-type calcium current (I_T) enables post-inhibitory rebound bursting characteristic of STN neurons, the calcium-activated AHP current provides spike-frequency adaptation through intracellular calcium dynamics (τ_Ca = 120 ms, half-activation [Ca] = 15.0 µM), and the hyperpolarization-activated cation current (I_H) contributes to autonomous pacemaking. Activation gating time constants for I_Na and I_K are scaled by 0.75 to match observed firing frequencies. Spikes are detected as upward crossings of 0 mV with a minimum inter-spike interval of 2 ms.

**Table 1a.**
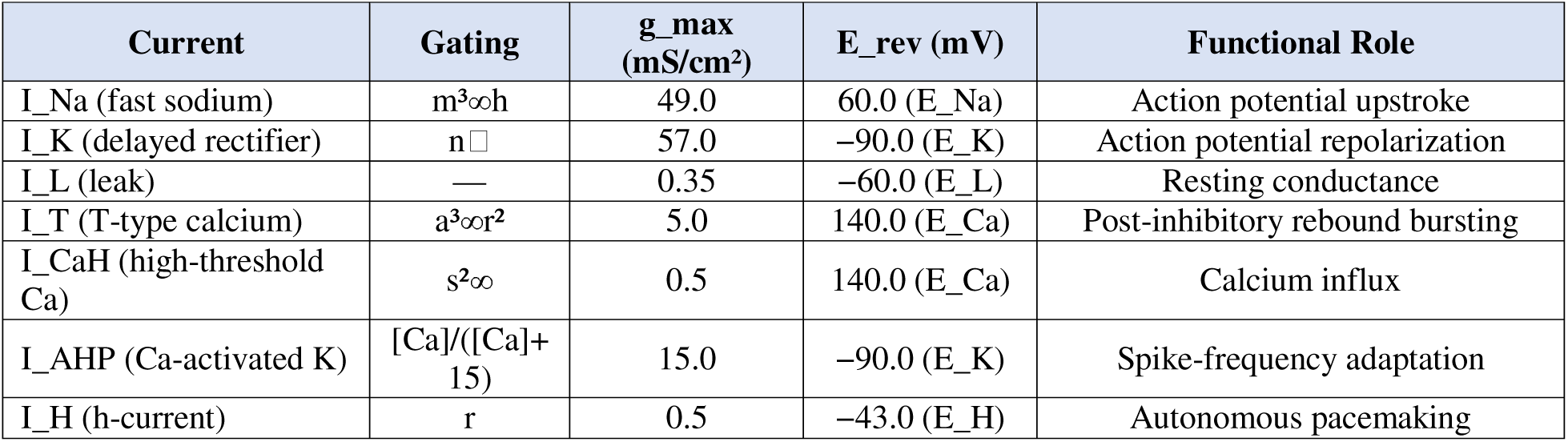
STN ionic current parameters (Gillies-Willshaw).

| Current | Gating | $g_{max}$<br>(mS/cm<sup>2</sup>) | $E_{rev}$ (mV) | Functional Role |
| --- | --- | --- | --- | --- |
| $I_{Na}$ (fast sodium) | $m^3\infty h$ | 49.0 | 60.0 ($E_{Na}$) | Action potential upstroke |
| $I_K$ (delayed rectifier) | $n^4$ | 57.0 | -90.0 ($E_K$) | Action potential repolarization |
| $I_L$ (leak) | — | 0.35 | -60.0 ($E_L$) | Resting conductance |
| $I_T$ (T-type calcium) | $a^3\infty r^2$ | 5.0 | 140.0 ($E_{Ca}$) | Post-inhibitory rebound bursting |
| $I_{CaH}$ (high-threshold Ca) | $s^2\infty$ | 0.5 | 140.0 ($E_{Ca}$) | Calcium influx |
| $I_{AHP}$ (Ca-activated K) | $[Ca]/([Ca]+15)$ | 15.0 | -90.0 ($E_K$) | Spike-frequency adaptation |
| $I_H$ (h-current) | $r$ | 0.5 | -43.0 ($E_H$) | Autonomous pacemaking |

#### 2.2.2 GPe and GPi Neurons

GPe and GPi neurons are modeled using the Rubin-Terman formalism [16], which provides the T-type calcium and calcium-activated potassium currents necessary for rebound bursting dynamics implicated in STN-GPe oscillatory interactions. The membrane equation follows the same Hodgkin-Huxley form with six ionic currents (Table 2b). GPe and GPi differ in their T-type calcium conductance, AHP conductance, and baseline applied current (I_app: GPe = 1.5 µA/cm², GPi = 2.0 µA/cm² at default, optimized as free parameters). Intracellular calcium decays with τ_Ca = 20 ms and gates the AHP current with half-activation [Ca] = 10. Spikes are detected at −20 mV threshold crossings. Gating variables are clamped to [0, 1] and voltages to [−100, 60] mV to ensure numerical stability.

**Table 1b.**
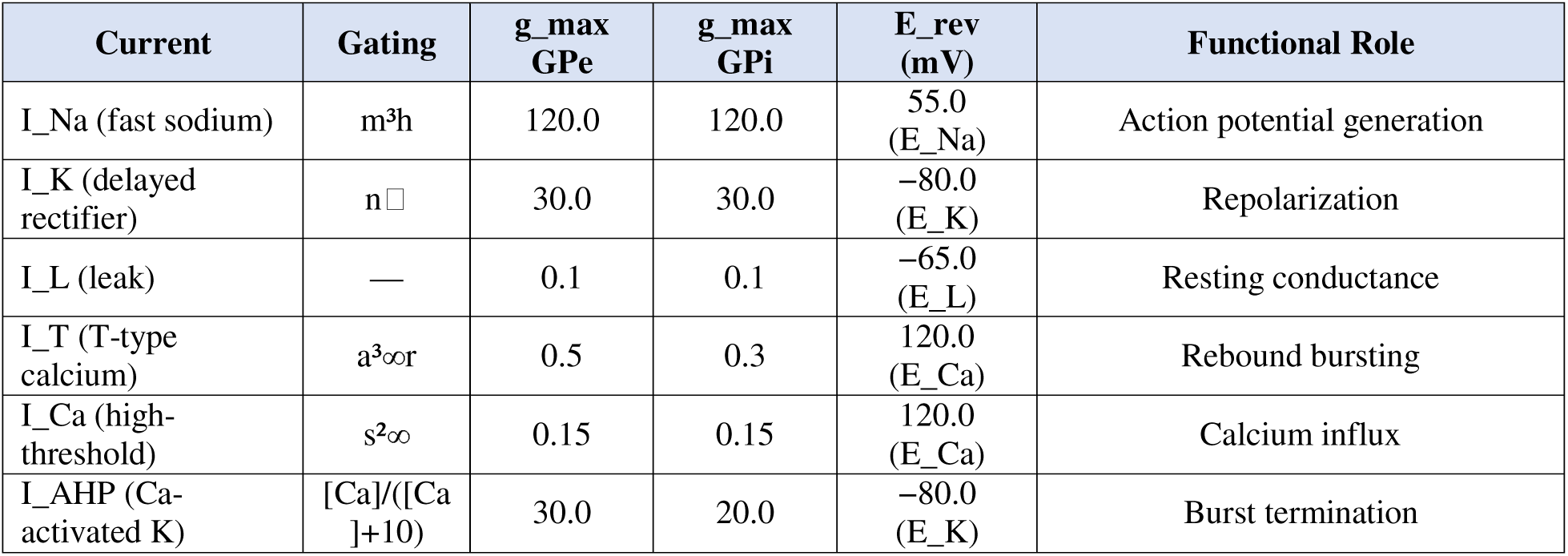
GPe/GPi ionic current parameters (Rubin-Terman).

| Current | Gating | $g_{max}$<br>GPe | $g_{max}$<br>GPi | $E_{rev}$<br>(mV) | Functional Role |
| --- | --- | --- | --- | --- | --- |
| $I_{Na}$ (fast sodium) | $m^3h$ | 120.0 | 120.0 | 55.0<br>($E_{Na}$) | Action potential generation |
| $I_K$ (delayed rectifier) | $n^4$ | 30.0 | 30.0 | $-80.0$<br>($E_K$) | Repolarization |
| $I_L$ (leak) | — | 0.1 | 0.1 | $-65.0$<br>($E_L$) | Resting conductance |
| $I_T$ (T-type calcium) | $a^3\infty r$ | 0.5 | 0.3 | 120.0<br>($E_{Ca}$) | Rebound bursting |
| $I_{Ca}$ (high-threshold) | $s^2\infty$ | 0.15 | 0.15 | 120.0<br>($E_{Ca}$) | Calcium influx |
| $I_{AHP}$ (Ca-activated K) | $[Ca]/([Ca] + 10)$ | 30.0 | 20.0 | $-80.0$<br>($E_K$) | Burst termination |

#### 2.2.3 Numerical Integration

All differential equations are integrated using the forward Euler method with a timestep of dt = 0.025 ms. This timestep was validated by comparison with dt = 0.0125 ms at 45,000 neurons: firing rates differed by less than 2 Hz across all populations, and the qualitative pattern of beta oscillation emergence was preserved.

### 2.3 Synaptic Transmission

Synaptic currents are computed using a double-exponential conductance-based model. For each synapse, the current is *I*_syn,*i* = *g*_max · *s*_ij · (*E*_syn − *V*_post,*i*), where *g*_max is peak conductance, *E*_syn is reversal potential, and *s*_ij is a gating variable with separate rise and decay time constants. Each presynaptic spike produces a conductance waveform through a double-exponential kernel. Total synaptic current onto each postsynaptic neuron is the sum over all presynaptic connections. Synaptic parameters are listed in Table 1. STN→GPe and STN→GPi synapses are excitatory (AMPA, *E*_syn = 0 mV, τ_rise = 5 ms, τ_decay = 3 ms). GPe→STN and GPe→GPi synapses are inhibitory (GABA_A, *E*_syn = −70 mV, τ_rise = 8/5 ms, τ_decay = 8 ms). Synaptic delays: 5 ms for STN→GPe, STN→GPi, and GPe→GPi; 8 ms for GPe→STN.

An asymmetric current scaling is applied in the network integrator to account for different unit conventions between neuron models. Synaptic currents arriving at STN neurons are passed directly. Synaptic currents arriving at GPe and GPi neurons are scaled by a factor of 0.005, and noise currents by 0.01. These scale factors were calibrated during initial network tuning and held constant throughout all experiments.

**Table 2.** Synaptic connectivity and parameters.

| Pathway | Type | K | $g_{\text{max}}$<br>(mS/cm<sup>2</sup>) | $\tau_{\text{rise}}$<br>(ms) | $\tau_{\text{decay}}$<br>(ms) | $E_{\text{syn}}$<br>(mV) | Delay<br>(ms) |
| --- | --- | --- | --- | --- | --- | --- | --- |
| STN→GPe | Excitatory<br>(AMPA) | 15 | 0.2 | 5.0 | 3.0 | 0 | 5.0 |
| GPe→STN | Inhibitory<br>(GABA) | 14 | 0.9 | 8.0 | 8.0 | -70 | 8.0 |
| STN→GPi | Excitatory<br>(AMPA) | 30 | 0.2 | 5.0 | 3.0 | 0 | 5.0 |
| GPe→GPi | Inhibitory<br>(GABA) | 10 | 0.3 | 5.0 | 8.0 | -70 | 5.0 |

### 2.4 Background Input

Each neuron receives an independent background current generated by an Ornstein-Uhlenbeck (OU) process, providing stochastic input that represents unmodeled afferent activity. The OU process has mean µ, standard deviation σ, and correlation time constant τ = 5 ms for all populations. Integration uses the Euler-Maruyama method. For STN neurons, the OU output is added directly as external current. For GPe and GPi neurons, the OU output is scaled by the noise scale factor (0.01) in the network integrator. The mean and standard deviation of the OU process for each population are free parameters in the optimization.

### 2.5 GPU-Accelerated Implementation

The network model is implemented in JAX [20], a numerical computing framework that transforms Python/NumPy code into optimized GPU kernels via the XLA compiler. Three features of JAX are central to achieving high performance. First, the implementation follows a purely functional programming paradigm in which all neuron and synapse state is represented as immutable arrays. Second, *jax.vmap* (vectorized map) transforms single-neuron step functions into population-level operations, eliminating Python-level loops. Third, the entire simulation loop is compiled into a single fused kernel using *jax.lax.scan*, which replaces the Python *for* loop over timesteps with a compiled iteration executing entirely on the GPU. Synaptic connectivity is stored in sparse coordinate (COO) format with scatter-add operations for postsynaptic current summation, reducing memory requirements from approximately 6.8 GB (dense) to under 100 MB (sparse) at 45,000 neurons. The computational workflow is illustrated in Figure 1.

**Figure 1:**
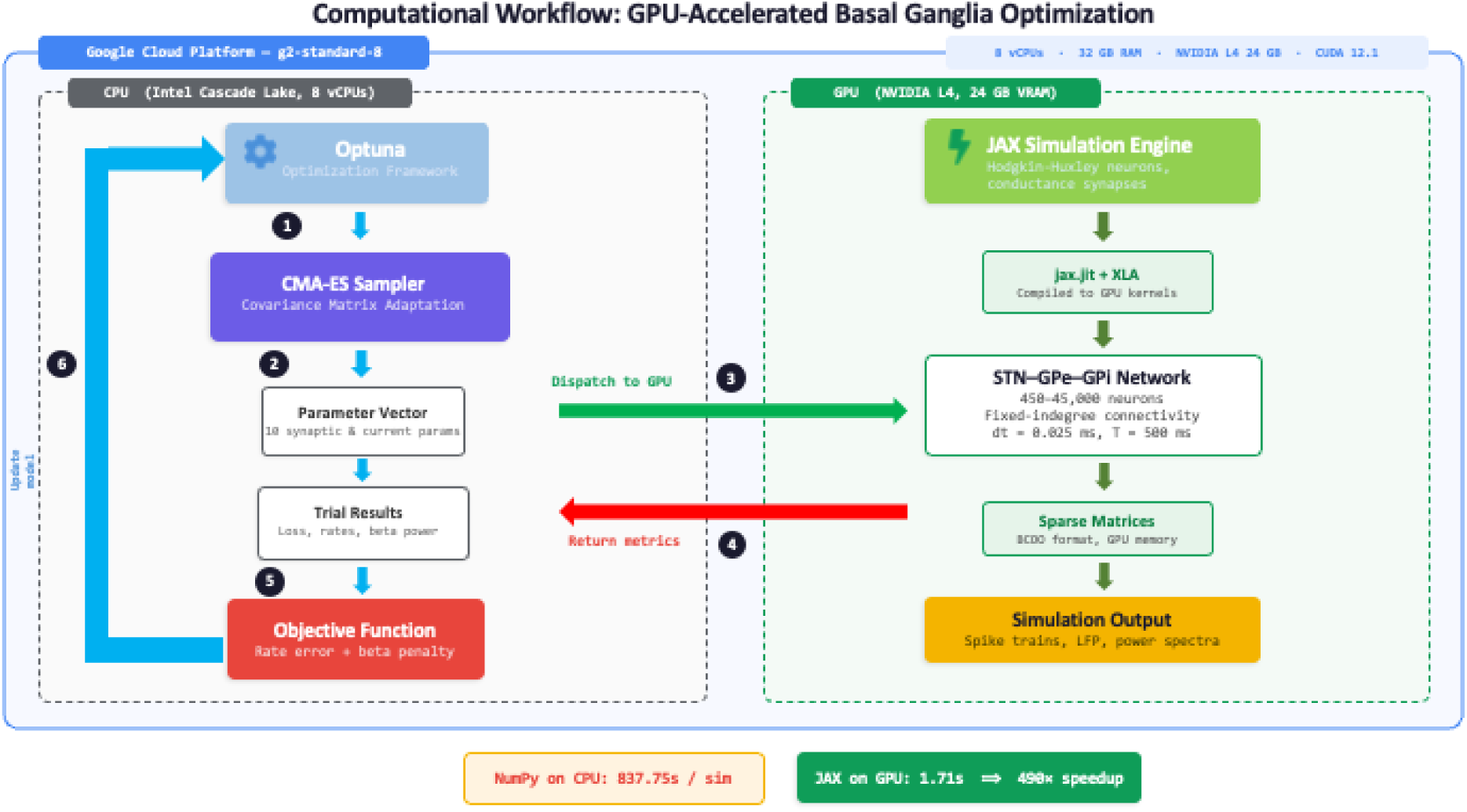
Computational workflow illustrating the GPU-accelerated pipeline. The CPU (Intel Cascade Lake, 8 vCPUs) runs the Optuna/CMA-ES optimizer, which dispatches 10-dimensional parameter vectors to the GPU (NVIDIA L4, 24 GB VRAM). The JAX simulation engine, compiled via XLA, executes the full network simulation and returns spike trains, LFP, and power spectra for metric computation. NumPy baseline: 837.75 s/sim; JAX on GPU: 1.71 s/sim (490× speedup).

### 2.6 Cloud Computing Infrastructure

All simulations were performed on Google Cloud Platform (GCP) using a g2-standard-8 virtual machine instance (us-central1-c). The instance provides 8 vCPUs (Intel Cascade Lake), 32 GB RAM, and a single NVIDIA L4 GPU with 24 GB VRAM running CUDA 12.1. The on-demand cost is approximately $0.84/hour, and a complete 1,000-trial optimization completes in under one hour (∼$2.50 total for all simulations). No local GPU hardware is required. All code and optimized parameters are available at https://github.com/neuronlab-cell/STN_GPe_GPi_Beta (Figure 2).

**Figure 2.**
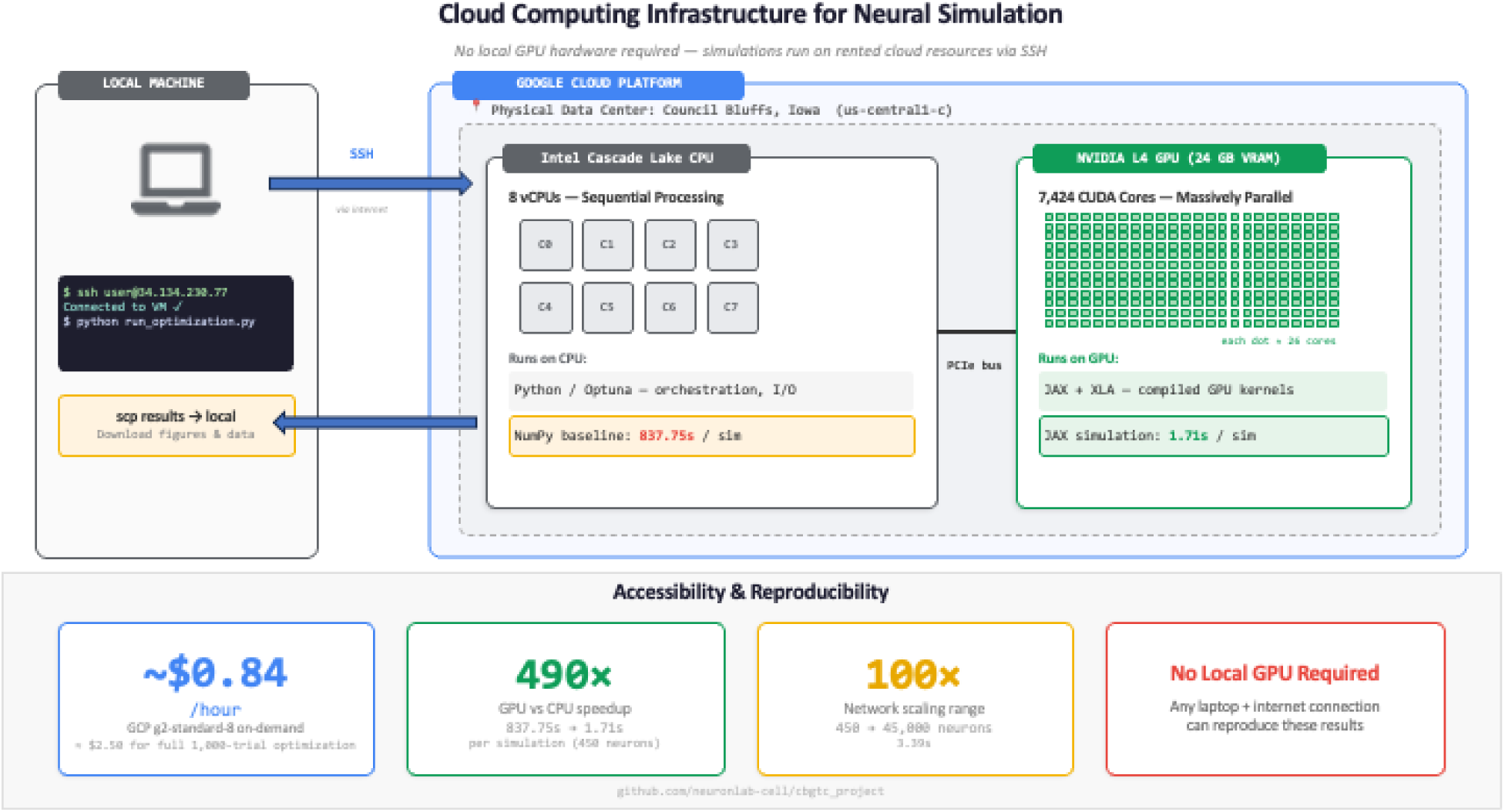
Cloud computing infrastructure. The entire pipeline runs on a Google Cloud Platform g2-standard-8 instance connected via SSH from any local machine. Key accessibility metrics: ∼$0.84/hour, 490× GPU vs CPU speedup, 100× network scaling range (450 to 45,000 neurons), no local GPU hardware required.

### 2.7 Optimization Procedure

Parameter optimization was performed using CMA-ES [21] within the Optuna framework [22]. CMA-ES was selected based on a sampler comparison study (5 seeds, 500 trials each) in which it achieved lower final loss than TPE (0.131 ± 0.013) and random sampling (0.257 ± 0.061), achieving 0.114 ± 0.007 (mean ± SD). Two optimization studies were conducted, both using 450 neurons with 400 ms simulations (dt = 0.025 ms) and 100 ms burn-in. Each optimized 10 parameters: six intrinsic parameters (*I*_STN, *I*_GPe, *I*_GPi, σ_STN, σ_GPe, σ_GPi) and four synaptic conductance multipliers scaling the baseline values in Table 2.

#### Study 1: Healthy State

500 CMA-ES trials with firing rate targets from the primate literature: STN ∼20 Hz [11, 3], GPe ∼70 Hz [13], GPi ∼80 Hz [13]. All four synaptic multipliers searched over [0.5, 2.0]. The objective function was *L*_healthy = 5.0 × Σ(normalized rate error)² + 0.2 × Σ(CV error)² + 15.0 × (beta penalty)².

#### Study 2: Parkinsonian State

1,000 CMA-ES trials with targets reflecting dopamine depletion: STN ∼27.5 Hz, GPe ∼42.5 Hz, GPi ∼82.5 Hz [11–13]. The search ranges for synaptic multipliers were set asymmetrically to reflect known directions of dopamine-dependent plasticity [25]: *g*_STN→GPe□[1.5, 5.0], *g*_GPe→STN□ [0.1, 0.8], *g*_STN→GPi□ [1.0, 4.0], *g*_GPe→GPi [0.2, 1.2]. These constraints encode the prior expectation that dopamine loss increases excitatory drive from STN and reduces inhibitory control from GPe, consistent with the classical rate model [25] and experimental observations [11, 12]. Within these bounds, the optimizer determined the specific magnitude of each change. The beta term penalized insufficient beta power (target: 20%), with an additional penalty above 40%.

### 2.8 Metrics and Analysis

Firing rates were computed as total spike count divided by neuron count and post-burn-in duration. CV of inter-spike intervals was computed per neuron and averaged across each population. A local field potential (LFP) proxy was computed as mean membrane voltage across a random subsample of 500 neurons (or all neurons if fewer available). Power spectral density was estimated using Welch’s method (segment length = min(signal length, 8,192 samples), Hann window, 50% overlap). Beta power fraction was defined as integrated power in 13–30 Hz divided by total power in 1–100 Hz, after DC removal. Spikes were detected as upward threshold crossings (0 mV for STN, −20 mV for GPe/GPi) with a 2 ms minimum ISI for STN.

### 2.9 Validation Pipeline

Four validation analyses were performed. *Scale invariance:* optimized parameters were applied without modification to networks of 450, 1,800, 4,500, 15,000, and 45,000 neurons (600 ms, 100 ms burn-in). *Statistical validation:* 10 random seeds at 45,000 neurons with independent t-tests. *Timestep sensitivity:* dt = 0.025 versus 0.0125 ms at 45,000 neurons. *Sampler comparison:* CMA-ES versus TPE versus random sampling (5 seeds × 500 trials, healthy objective).

### 2.10 DBS Simulation

DBS was modeled as an informational lesion [24] via two parameter modifications to the Parkinsonian configuration: increased STN tonic drive (*I*_STN: 70.0 → 150.0 µA/cm²) and further reduction of GPe→STN coupling (*g*_GPe→STN multiplier: 0.11 → 0.05). This follows the hypothesis that DBS desynchronizes pathological activity by overriding intrinsic STN dynamics [16, 24]. The simulation was run at 45,000 neurons (600 ms, 100 ms burn-in).

## 3. Results

### 3.1 GPU Acceleration Enables High-Throughput Parameter Optimization

The JAX-based implementation achieved a 490-fold speedup over the conventional NumPy/CPU baseline. A 600 ms simulation of the 450-neuron network completed in 1.71 seconds on the NVIDIA L4 GPU, compared to 837.75 seconds using a Python-loop implementation on the same instance’s CPU (Table 2). This acceleration was the prerequisite for systematic parameter optimization: a 1,000-trial CMA-ES optimization that would require approximately 233 hours on CPU completed in under one hour on GPU.

Performance scaled favorably with network size. Simulation wall-clock time increased from 1.71 seconds at 450 neurons to only 3.39 seconds at 45,000 neurons, which represents a 100-fold increase in network size for less than a 2-fold increase in computation time (Table 2). Among optimization samplers, CMA-ES consistently achieved the lowest final loss (0.114 ± 0.007), outperforming TPE (0.131 ± 0.013) and random sampling (0.257 ± 0.061). The complete workflow (Figure 1) and infrastructure (Figure 2) demonstrate that the pipeline executes on a single commodity cloud instance (∼$0.84/hour).

**Table 2.**
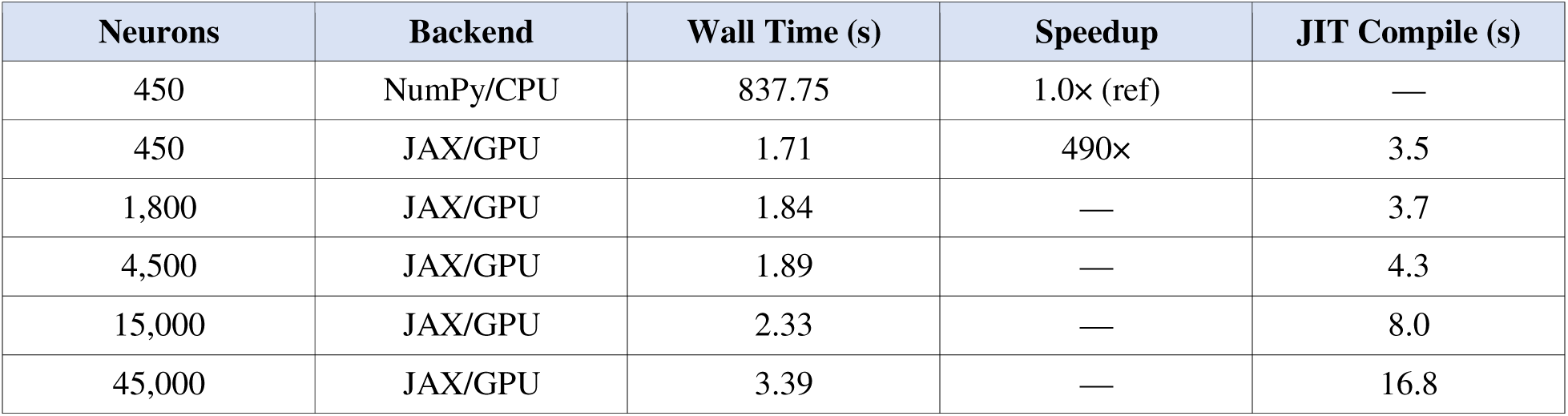
Performance benchmarks (600 ms simulation).

| Neurons | Backend | Wall Time (s) | Speedup | JIT Compile (s) |
| --- | --- | --- | --- | --- |
| 450 | NumPy/CPU | 837.75 | 1.0× (ref) | — |
| 450 | JAX/GPU | 1.71 | 490× | 3.5 |
| 1,800 | JAX/GPU | 1.84 | — | 3.7 |
| 4,500 | JAX/GPU | 1.89 | — | 4.3 |
| 15,000 | JAX/GPU | 2.33 | — | 8.0 |
| 45,000 | JAX/GPU | 3.39 | — | 16.8 |

### 3.2 Healthy Network Configuration Reproduces Normative Basal Ganglia Activity

Optimization of the healthy network (500 CMA-ES trials, 10 parameters) yielded a configuration that closely approximated literature-derived firing rate targets [3, 11, 13]. STN neurons fired at 26.4 Hz (target: 20 Hz), GPe at 62.3 Hz (target: 70 Hz), and GPi at 82.3 Hz (target: 80 Hz). The healthy network exhibited asynchronous, irregular spiking activity (Figure 3, top row) with no evidence of population-wide synchronization. The STN LFP proxy displayed low-amplitude, aperiodic fluctuations (Figure 4, top panel), and spectral analysis confirmed minimal beta-band content: only 0.4% of total STN spectral power fell within the 13–30 Hz range (Figure 5A, blue trace; Figure 6B).

**Figure 3.**
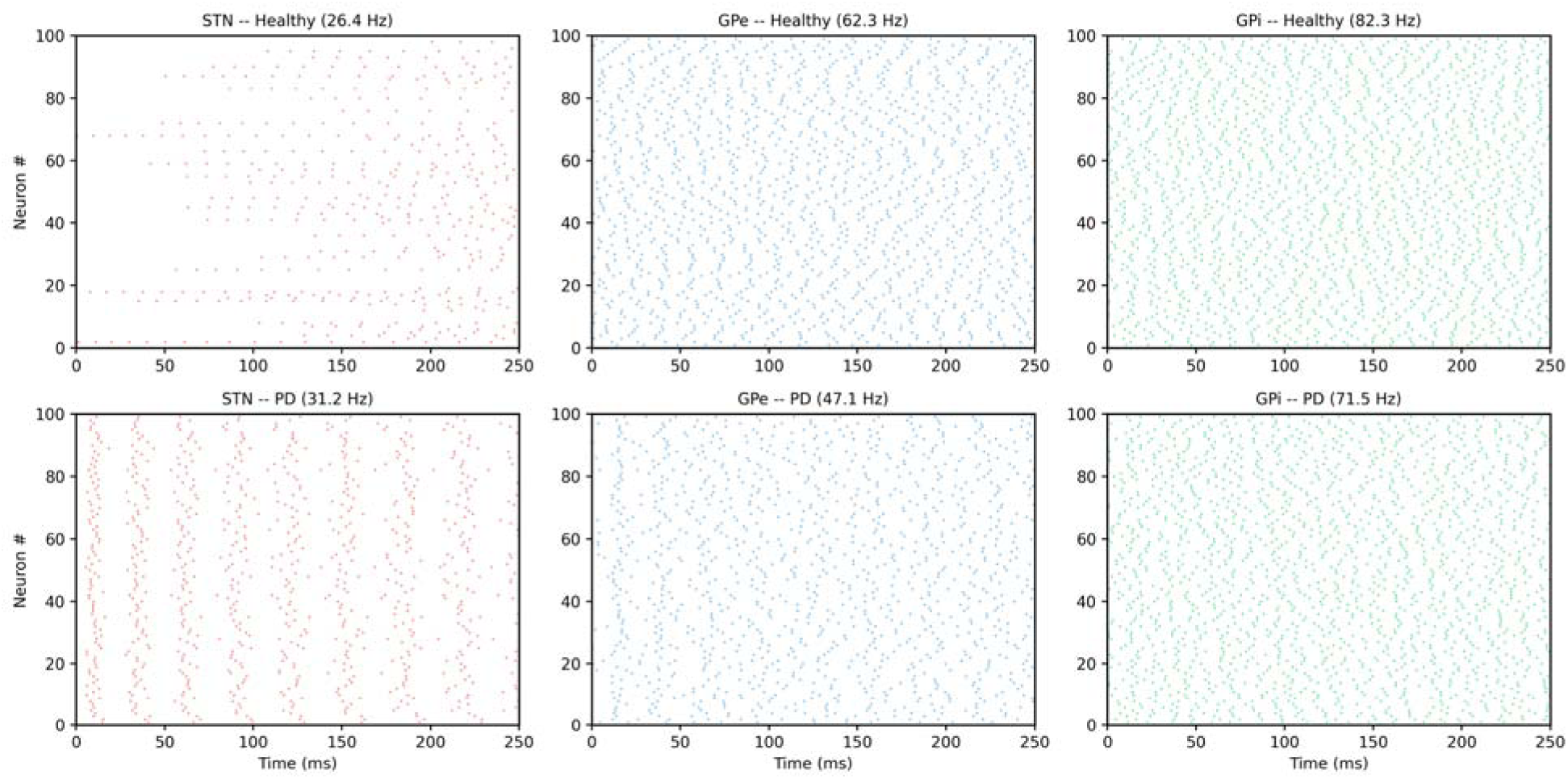
Raster plots showing spiking activity of STN, GPe, and GPi populations in healthy (top row) and Parkinsonian (bottom row) conditions. 100 neurons per population are displayed. Mean firing rates are indicated above each panel. Note the emergence of synchronized bursting in the Parkinsonian STN at approximately 20 Hz.

**Figure 4.**
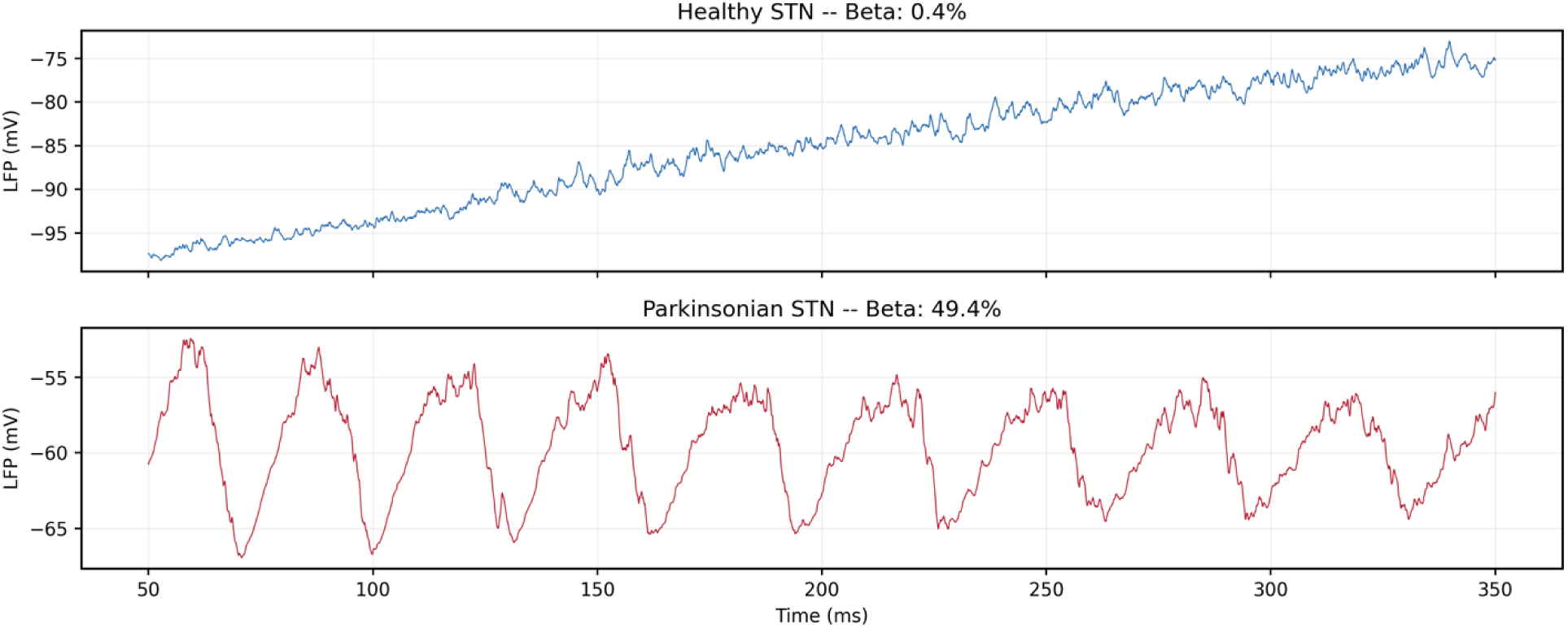
STN local field potential (LFP) proxy traces. Top: healthy condition showing low-amplitude aperiodic fluctuations (beta: 0.4%). Bottom: Parkinsonian condition showing prominent rhythmic oscillations at approximately 20 Hz (beta: 49.4%).

**Figure 5.**
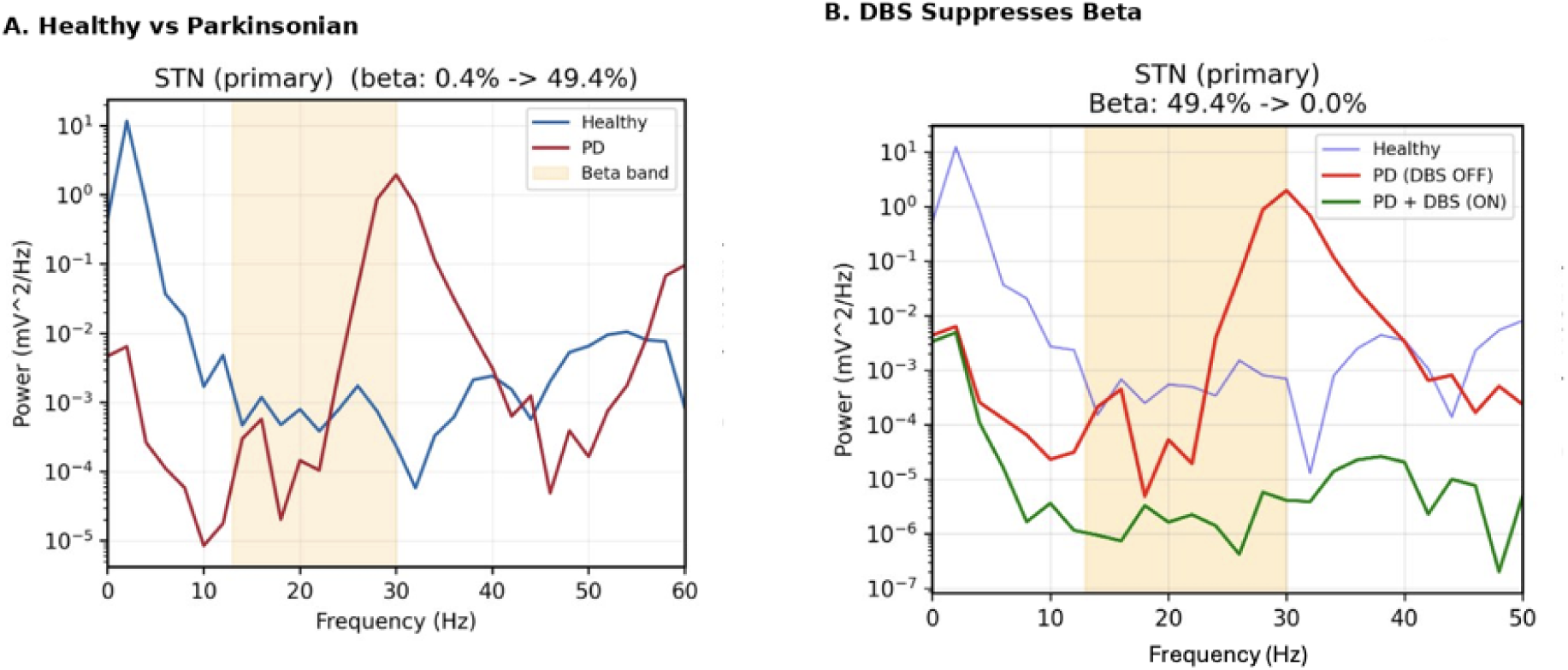
Power spectral density of the STN local field potential proxy. (A) Healthy (blue) versus Parkinsonian (red) conditions. The shaded region indicates the beta band (13–30 Hz). Beta power fraction increases from 0.4% to 49.4%. (B) Effect of simulated DBS on the Parkinsonian network. DBS (green) eliminates the beta-band peak, reducing beta power from 49.4% to 0.0%.

**Figure 6.**
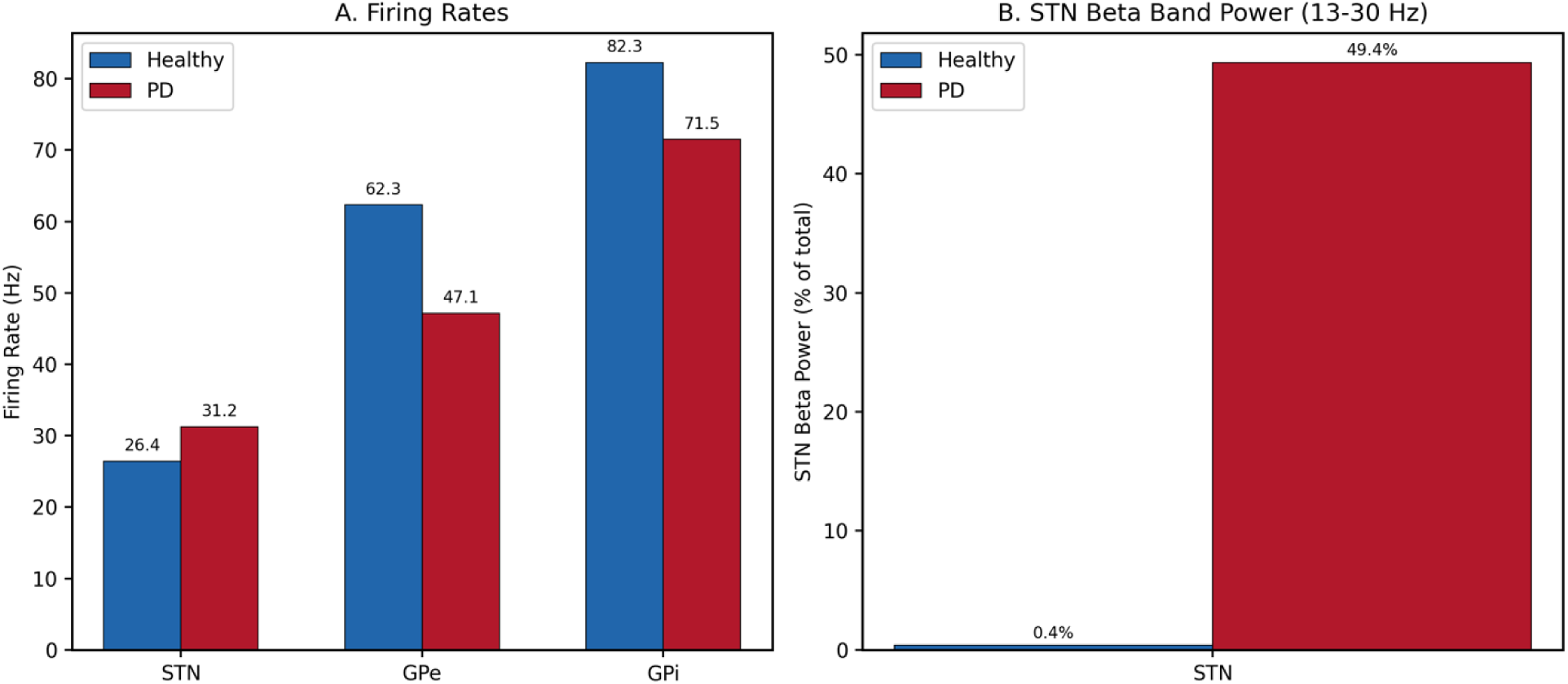
Summary of population-level metrics. (A) Mean firing rates for STN, GPe, and GPi in healthy (blue) and Parkinsonian (red) conditions. (B) STN beta power fraction in healthy versus Parkinsonian conditions.

### 3.3 Optimization Reveals Synaptic Reorganization in the Parkinsonian State

Optimization of the Parkinsonian network (1,000 CMA-ES trials with biologically motivated search bounds; see Section 2.7) converged on a configuration characterized by coordinated changes across multiple synaptic pathways. Comparison of the Parkinsonian and healthy synaptic multipliers revealed a distinctive pattern (Figure 7; Table 3): STN→GPe excitation increased 2.21-fold (healthy: 1.87×, PD: 4.13×); GPe→STN inhibition collapsed to 0.11× its healthy value (healthy: 0.99×, PD: 0.11×), representing an 89% reduction; STN→GPi excitation remained near baseline (PD/Healthy: 1.06×); and GPe→GPi inhibition increased moderately (PD/Healthy: 1.45×).

**Figure 7.**
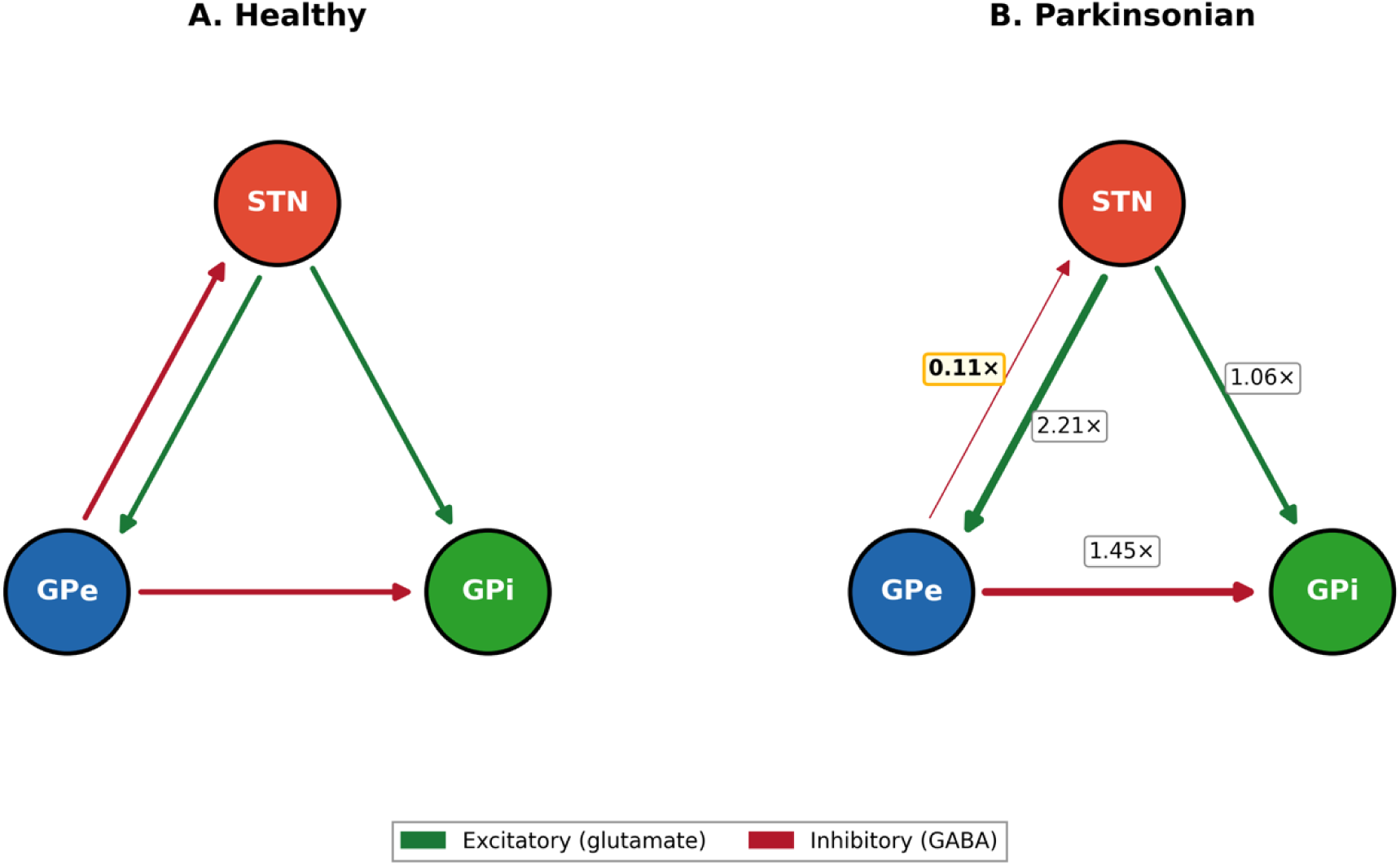
Network schematic showing synaptic organization in healthy (A) and Parkinsonian (B) states. Green arrows indicate excitatory (glutamatergic) projections, red arrows indicate inhibitory (GABAergic) projections. PD/Healthy synaptic multiplier ratios are shown: STN→GPe 2.21×, GPe→STN 0.11×, STN→GPi 1.06×, GPe→GPi 1.45×.

**Table 3.** Optimized parameters and PD/Healthy synaptic ratios.

| Parameter | Healthy | Parkinsonian | PD/Healthy | Direction |
| --- | --- | --- | --- | --- |
| I_STN ($\mu\text{A}/\text{cm}^2$) | 132.2 | 70.0 | — | ↓ Decreased |
| I_GPe ($\mu\text{A}/\text{cm}^2$) | 3.04 | 1.66 | — | ↓ Decreased |
| I_GPi ($\mu\text{A}/\text{cm}^2$) | 2.21 | 1.67 | — | ↓ Decreased |
| $\sigma_{\text{STN}}$ | 3.36 | 1.97 | — | ↓ Decreased |
| $\sigma_{\text{GPe}}$ | 33.4 | 66.4 | — | ↑ Increased |
| $\sigma_{\text{GPi}}$ | 67.3 | 96.4 | — | ↑ Increased |
| g_STN→GPe mult | 1.87 | 4.13 | **2.21×** | ↑ Increased |
| g_GPe→STN mult | 1.00 | 0.11 | **0.11×** | ↓↓ Collapsed |
| g_STN→GPi mult | 1.83 | 1.94 | **1.06×** | ≈ Unchanged |
| g_GPe→GPi mult | 0.69 | 1.00 | **1.45×** | ↑ Moderate |

The net effect is an asymmetric destabilization of the STN-GPe reciprocal loop: excitatory drive from STN to GPe escalates while the inhibitory return pathway from GPe to STN fails. This removes the negative feedback that normally stabilizes STN activity, creating conditions favorable for self-sustaining oscillatory dynamics.

### 3.4 Parkinsonian Configuration Produces Robust Beta-Band Oscillations

The Parkinsonian network exhibited striking differences from the healthy state. Firing rates shifted in directions consistent with experimental observations in MPTP-treated primates [11, 12]: STN rate increased from 26.4 to 31.2 Hz, GPe rate decreased from 62.3 to 47.1 Hz, and GPi rate decreased from 82.3 to 71.5 Hz (Figure 6A). STN beta power fraction increased 124x from 0.4% to 49.4% (Figure 5A; Figure 6B). This oscillatory activity was visible as synchronized population bursting in STN raster plots at approximately 20 Hz (Figure 3, bottom row). The STN LFP proxy showed clear rhythmic fluctuations contrasting with the aperiodic healthy LFP (Figure 4). The beta power fraction achieved (49.4%) exceeded the 20–40% target range, but remains within the range reported in severely dopamine-depleted conditions [2, 4].

### 3.5 Network Dynamics Are Invariant Across a 100-Fold Range of Network Sizes

Both parameter sets were applied without modification to networks from 450 to 45,000 neurons (Table 4; Figure 8). Healthy STN rates varied between 25.1 and 27.5 Hz (maximum deviation: <2.4 Hz), while Parkinsonian STN rates varied between 31.1 and 31.4 Hz (<0.3 Hz). Beta power in the Parkinsonian STN remained consistently elevated at approximately 50% (range: 45.7–51.2%) while healthy beta stayed below 5% at all sizes. This scale invariance is a direct consequence of the fixed-indegree connectivity scheme [23]. Simulation wall-clock time scaled sub-linearly, from 1.7 s at 450 neurons to 3.4 s at 45,000 neurons (Figure 9C).

**Figure 8.**
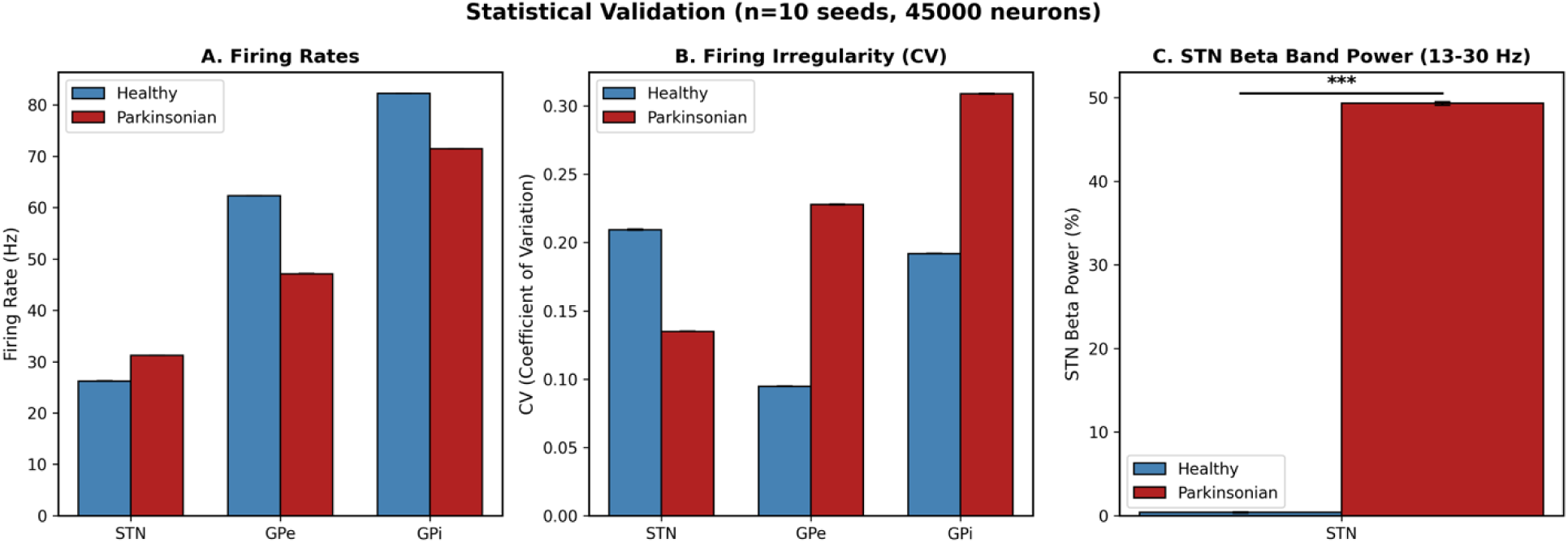
Statistical validation across 10 random seeds at 45,000 neurons. (A) Firing rates. (B) Coefficient of variation of inter-spike intervals. (C) STN beta power fraction. Error bars represent standard deviation. *** indicates p < 0.001 (independent t-test).

**Figure 9.**
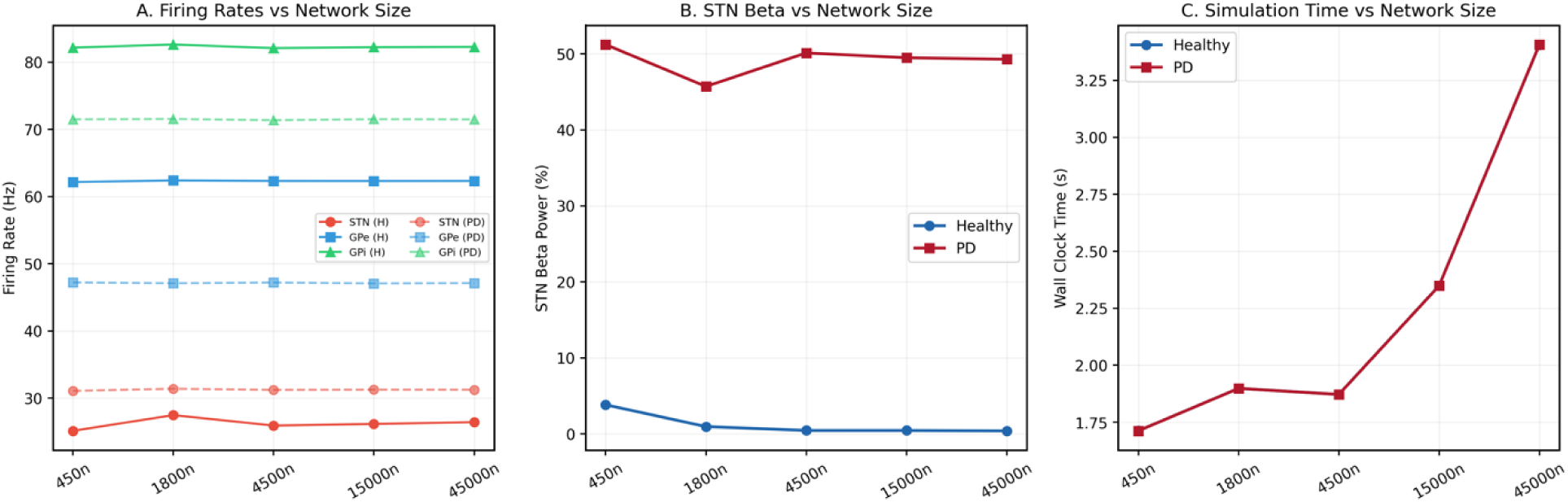
Scale invariance of network dynamics across 450 to 45,000 neurons. (A) Firing rates remain stable (<2.4 Hz deviation) across all sizes for both conditions. (B) STN beta power fraction is consistently elevated (∼50%) in the Parkinsonian condition and suppressed (<5%) in the healthy condition. (C) GPU simulation wall-clock time scales sub-linearly with network size.

**Table 4.**
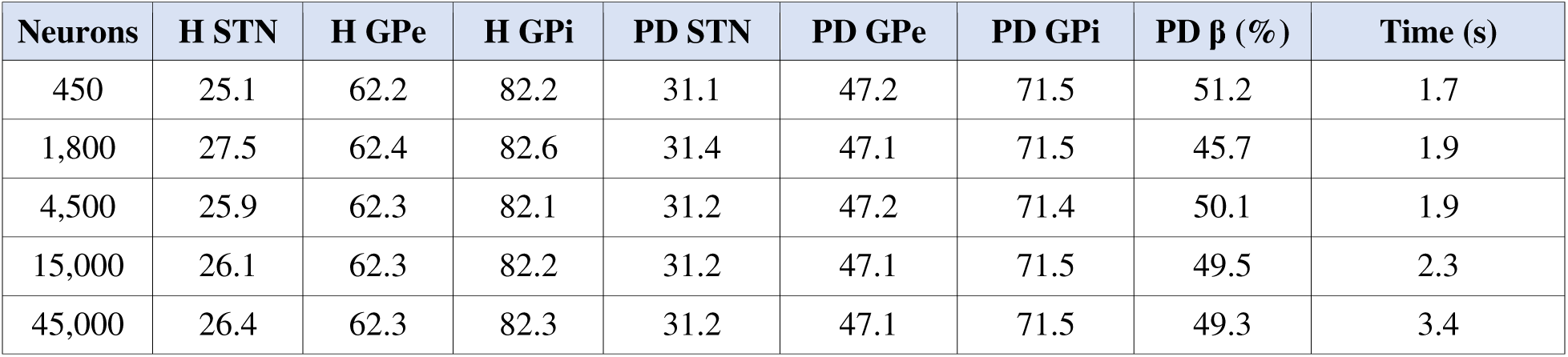
Scale invariance: firing rates and beta power across network sizes.

### 3.6 Statistical Validation Confirms Robustness Across Random Seeds

Both simulations were repeated with 10 different random seeds at 45,000 neurons (Figure 9; Table 5). All electrophysiological metrics showed highly significant differences between conditions (independent t-tests, all *p* < 0.001). Within-condition variability was extremely low: STN firing rate SD was 0.06 Hz (healthy) and 0.01 Hz (Parkinsonian). STN beta power fraction was 0.38 ± 0.06% (healthy) versus 49.33 ± 0.21% (Parkinsonian). The low variance indicates that the model’s behavior is determined primarily by parameters rather than stochastic initial conditions.

**Table 5.** Statistical validation (n = 10 seeds, 45,000 neurons).

| Metric | Healthy (mean ± SD) | Parkinsonian (mean ± SD) | p-value |
| --- | --- | --- | --- |
| STN Rate (Hz) | 26.23 ± 0.06 | 31.23 ± 0.01 | < 0.001 |
| GPe Rate (Hz) | 62.32 ± 0.01 | 47.12 ± 0.02 | < 0.001 |
| GPi Rate (Hz) | 82.24 ± 0.03 | 71.47 ± 0.02 | < 0.001 |
| STN CV | 0.21 ± 0.01 | 0.13 ± 0.01 | < 0.001 |
| GPe CV | 0.08 ± 0.01 | 0.23 ± 0.01 | < 0.001 |
| GPi CV | 0.19 ± 0.01 | 0.32 ± 0.01 | < 0.001 |
| STN Beta (%) | 0.38 ± 0.06 | 49.33 ± 0.21 | < 0.001 |

### 3.7 Simulated DBS Suppresses Pathological Beta Oscillations

DBS produced complete suppression of pathological beta oscillations (Table 6; Figure 5B). STN beta power fraction dropped from 49.4% to 0.0%. GPi firing rate recovered from 71.5 Hz (Parkinsonian) to 82.2 Hz, closely matching the healthy value (82.3 Hz). STN firing rate increased to 73.2 Hz, reflecting the elevated tonic drive. This elevated rate, while non-physiological, is consistent with the informational lesion interpretation: DBS overrides pathological STN dynamics with desynchronized high-frequency tonic activity that no longer drives beta oscillations in the downstream circuit [16, 24].

**Table 6.**
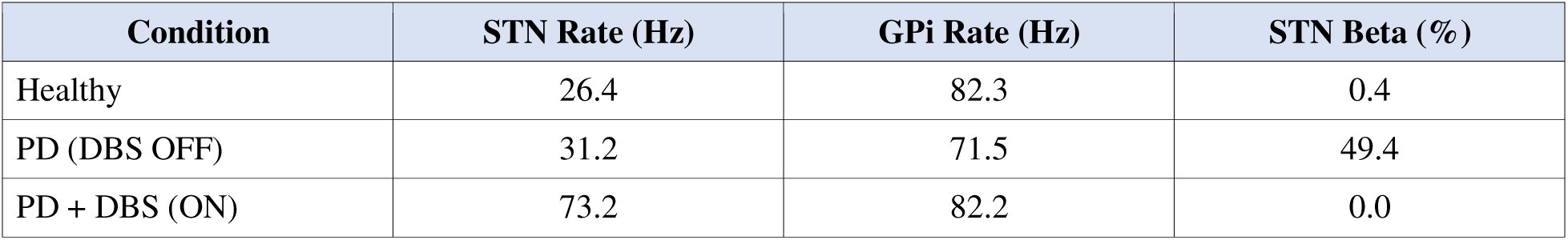
DBS simulation results (45,000 neurons).

## 4. Discussion

### 4.1 Biological Interpretation: Asymmetric Destabilization of the STN-GPe Loop

The central biological finding of this study is that systematic optimization, guided by experimentally derived firing rate and oscillatory targets, converges on a specific pattern of synaptic reorganization in the Parkinsonian state. Within biologically motivated search bounds reflecting known directions of dopamine-dependent plasticity, the optimizer identified a configuration dominated by two asymmetric changes within the STN-GPe reciprocal loop: a 2.21-fold increase in STN→GPe excitation and an 89% collapse of GPe→STN inhibition. This pattern was sufficient to transform an asynchronous, non-oscillatory network into one exhibiting robust beta-band oscillations at 49.4% of total spectral power.

This result is consistent with several lines of experimental evidence. Dopamine depletion strengthens glutamatergic transmission at subthalamic synapses [26, 27] while weakening pallidal inhibitory control [12, 28]. Our results suggest that the ratio of these changes may be more important than the absolute magnitude of any single change. This is compatible with the proposal by Plenz and Kital [15] that the STN-GPe network functions as a pacemaker circuit whose oscillatory properties depend critically on the balance of excitation and inhibition within the loop.

### 4.2 Methodological Contribution: Optimization-Driven Discovery

This work demonstrates a methodological approach in which high-performance computing enables systematic parameter space exploration as an alternative to manual tuning. Traditional basal ganglia models [16, 14, 17] typically establish parameter sets through iterative manual adjustment, which is time-consuming and difficult to reproduce. The GPU-accelerated workflow completes the pipeline end to end in under one hour at ∼$2.50 on commodity cloud hardware, making systematic exploration feasible as a routine modeling tool.

The two-stage optimization design mirrors the biological logic of disease modeling: the pathological state is defined relative to a normative reference. CMA-ES, which maintains a covariance model of the parameter space, was critical for navigating the 10-dimensional landscape efficiently (final loss 0.114 ± 0.007 versus 0.131 ± 0.013 for TPE and 0.257 ± 0.061 for random). The fixed-indegree connectivity scheme [23] ensured that parameters optimized at 450 neurons transferred to networks up to 100 times larger without retuning, providing confidence that the observed dynamics are genuine emergent properties rather than artifacts of a specific network realization.

### 4.3 Comparison with Existing Basal Ganglia Models

The present model builds on substantial prior computational work. Rubin and Terman [16] demonstrated that STN-GPe interactions can generate pathological rhythms using a 16-neuron network with manually tuned parameters. Hahn and McIntyre [19] extended this to larger networks but similarly relied on hand-tuning. Pavlides et al. [31] used mean-field approximations to analyze STN-GPe oscillatory regimes analytically. Our approach differs in three respects: systematic optimization replacing manual tuning, GPU acceleration enabling network sizes approaching primate basal ganglia scale (45,000 neurons), and a reproducible, cloud-accessible pipeline. However, several existing models incorporate features we omit, including cortex-basal ganglia-thalamic loops [29, 32], multi-compartment neurons [33], and striatal projections [18].

### 4.4 Limitations

Several limitations should be considered. The Parkinsonian search ranges encoded prior knowledge about dopamine-dependent synaptic changes and were not unconstrained. The model simplifies cortical input, the striatum, and thalamic feedback as intrinsic current and noise parameters, however, cortical beta oscillations likely contribute to subcortical beta through cortico-subthalamic projections [29, 30]. All neurons are single-compartment, which cannot capture dendritic processing or morphological diversity. The DBS simulation uses an informational lesion approach rather than explicit extracellular stimulation with electrode geometry. Finally, the model predictions have not been directly validated against experimental measurements of synaptic strength changes in dopamine-depleted preparations.

### 4.5 Future Directions

The most immediate extension is incorporation of cortical input to examine interactions between cortically generated and subcortically generated beta rhythms. Phase-amplitude coupling between cortical gamma and subcortical beta has been identified as a potential biomarker for adaptive DBS [34], and a coupled cortex-basal ganglia model could investigate its circuit mechanisms. The optimization framework is extensible to additional synaptic pathways, multiple neuron types, and neuromodulatory effects beyond dopamine. Finally, the simulation platform’s speed makes it suitable for generating synthetic training data for machine learning approaches to closed-loop DBS [35], supplementing limited clinical data currently available.

## Use of Generative AI

Generative AI tools were used to assist with improving the readability, formatting, and documentation of the manuscript and associated codebase. All AI-generated text was critically reviewed and edited by the authors. No generative AI tools were used to design the computational model, execute simulations, analyze data, or generate scientific conclusions.

## Data and Code Availability

All code, optimized parameters, and generated figures are publicly available at https://github.com/neuronlab-cell/STN_GPe_GPi_Beta. Optimization results and raw simulation data are provided in the repository’s results/ directory, enabling full reproduction of the results reported here on identical cloud infrastructure.

## Conflicts of Interest, Funding, and Ethics Statement

The authors have declared no competing interests exist. The funders had no role in study design, data collection and analysis, decision to publish, or preparation of the manuscript. Ethical approval was not required for this study.

## Author Approval

All authors have seen and approved the manuscript. This manuscript has not been accepted or published at any other venue.

## References

[1] GBD 2016 Parkinson’s Disease Collaborators. Global, regional, and national burden of Parkinson’s disease, 1990–2016. Lancet Neurol. 17, 939–953 (2018).

[2] Brown P. Oscillatory nature of human basal ganglia activity: relationship to the pathophysiology of Parkinson’s disease. Mov. Disord. 18, 357–363 (2003).

[3] Levy R, Hutchison W D, Lozano A M and Dostrovsky J O. High-frequency synchronization of neuronal activity in the subthalamic nucleus of Parkinsonian patients with limb tremor. J. Neurosci. 20, 7766–7775 (2000).

[4] Kühn A A, Kupsch A, Schneider G-H and Brown P. Reduction in subthalamic 8–35 Hz oscillatory activity correlates with clinical improvement in Parkinson’s disease. Eur. J. Neurosci. 23, 1956–1960 (2006).

[5] Neumann W-J et al. Subthalamic synchronized oscillatory activity correlates with motor impairment in patients with Parkinson’s disease. Mov. Disord. 31, 1748–1751 (2016).

[6] Weinberger M et al. Beta oscillatory activity in the subthalamic nucleus and its relation to dopaminergic response in Parkinson’s disease. J. Neurophysiol. 96, 3248–3256 (2006).

[7] Kühn A A et al. High-frequency stimulation of the subthalamic nucleus suppresses oscillatory β activity in patients with Parkinson’s disease in parallel with improvement in motor performance. J. Neurosci. 28, 6165–6173 (2008).

[8] Little S et al. Adaptive deep brain stimulation in advanced Parkinson disease. Ann. Neurol. 74, 449–457 (2013).

[9] Little S and Brown P. Debugging adaptive deep brain stimulation for Parkinson’s disease. Mov. Disord. 35, 555–561 (2020).

[10] Meidahl A C et al. Adaptive deep brain stimulation for movement disorders: the long road to clinical therapy. Mov. Disord. 32, 810–819 (2017).

[11] Bergman H, Wichmann T, Karmon B and DeLong M R. The primate subthalamic nucleus. II. Neuronal activity in the MPTP model of parkinsonism. J. Neurophysiol. 72, 507–520 (1994).

[12] Filion M and Tremblay L. Abnormal spontaneous activity of globus pallidus neurons in monkeys with MPTP-induced parkinsonism. Brain Res. 547, 142–151 (1991).

[13] DeLong M R. Activity of pallidal neurons during movement. J. Neurophysiol. 34, 414–427 (1971).

[14] Terman D, Rubin J E, Yew A C and Wilson C J. Activity patterns in a model of the subthalamopallidal network. J. Neurosci. 22, 2963–2976 (2002).

[15] Plenz D and Kital S T. A basal ganglia pacemaker formed by the subthalamic nucleus and external globus pallidus. Nature 400, 677–682 (1999).

[16] Rubin J E and Terman D. High frequency stimulation of the subthalamic nucleus eliminates pathological thalamic rhythmicity in a computational model. J. Comput. Neurosci. 16, 211–235 (2004).

[17] Gillies A and Willshaw D. Membrane channel interactions underlying rat subthalamic projection neuron rhythmic and bursting activity. J. Neurophysiol. 95, 2352–2365 (2006).

[18] Humphries M D, Stewart R D and Gurney K N. A physiologically plausible model of action selection and oscillatory activity in the basal ganglia. J. Neurosci. 26, 12921–12942 (2006).

[19] Hahn P J and McIntyre C C. Modeling shifts in the rate and pattern of subthalamopallidal network activity during deep brain stimulation. J. Comput. Neurosci. 28, 425–441 (2010).

[20] Bradbury J, et al. JAX: composable transformations of Python+NumPy programs. GitHub (2018). http://github.com/google/jax

[21] Hansen N. The CMA evolution strategy: a tutorial. arXiv:1604.00772 (2016).

[22] Akiba T, Sano S, Yanase T, Ohta T and Koyama M. Optuna: A next-generation hyperparameter optimization framework. Proc. 25th ACM SIGKDD 2623–2631 (2019).

[23] Gerstner W, Kistler W M, Naud R and Paninski L. Neuronal Dynamics: From Single Neurons to Networks and Models of Cognition (Cambridge University Press, 2014), Ch. 12.3.

[24] Grill W M, Snyder A N and Miocinovic S. Deep brain stimulation creates an informational lesion of the stimulated nucleus. NeuroReport 15, 1137–1140 (2004).

[25] Albin R L, Young A B and Penney J B. The functional anatomy of basal ganglia disorders. Trends Neurosci. 12, 366–375 (1989).

[26] Shen W, Flajolet M, Greengard P and Surmeier D J. Dichotomous dopaminergic control of striatal synaptic plasticity. Science 321, 848–851 (2008).

[27] Chu H-Y, McIver E L, Kovaleski R F, Atherton J F, Bevan M D. Loss of hyperdirect pathway cortico-subthalamic inputs following degeneration of midbrain dopamine neurons. Neuron 95, 1306–1318 (2017).

[28] Kita H and Kita T. Cortical stimulation evokes abnormal responses in the dopamine-depleted rat basal ganglia. J. Neurosci. 31, 10311–10322 (2011).

[29] Holgado A J N, Terry J R and Bogacz R. Conditions for the generation of beta oscillations in the subthalamic nucleus–globus pallidus network. J. Neurosci. 30, 12340–12352 (2010).

[30] Shimamoto S A et al. Subthalamic nucleus neurons are synchronized to primary motor cortex local field potentials in Parkinson’s disease. J. Neurosci. 33, 7220–7233 (2013).

[31] Pavlides A, Hogan S J and Bogacz R. Computational models describing possible mechanisms for generation of excessive beta oscillations in Parkinson’s disease. PLoS Comput. Biol. 11, e1004609 (2015).

[32] Kumaravelu K, Brocker D T and Grill W M. A biophysical model of cortical–basal ganglia–thalamic network in 6-OHDA lesioned rat model of Parkinson’s Disease. J. Comput. Neurosci. 40, 175–196 (2016).

[33] Gunay C, Edgerton J R and Jaegar D. Channel density distributions explain spiking variability in the globus pallidus. J. Neurosci. 28, 7476–7491 (2008).

[34] de Hemptinne C et al. Therapeutic deep brain stimulation reduces cortical phase-amplitude coupling in Parkinson’s disease. Nat. Neurosci. 18, 779–786 (2015).

[35] Gilron R et al. Long-term wireless streaming of neural recordings for circuit discovery and adaptive stimulation in individuals with Parkinson’s disease. Nat. Biotechnol. 39, 1078–1085 (2021).

